# Practice often and always get ready: a spiking mechanistic model for voluntary motor control

**DOI:** 10.1101/2023.06.02.543521

**Authors:** Chen Zhao, He Cui

**Affiliations:** CAS Center for Excellence in Brain Science and Intelligent Technology, Institute of Neuroscience, Shanghai, 200031, China; Shanghai Center for Brain and Brain-inspired Intelligence Technology, Shanghai, 200031, China

## Abstract

In many voluntary movement, neural activities ranging from cortex to spinal cord can be roughly described as the stages of motor intention, preparation, and execution. Recent advances in neuroscience have proposed many theories to understand how motor intention can be transformed into action following these stages, but they still lack a holistic and mechanistic theory to account for the whole process. Here, we try to formulate this question by abstracting two underlying principles: 1) the neural system is specializing the final motor command through a hierarchical network by multitudes of training supervised by the action feedback (“practice often”); 2) prediction is a general mechanism throughout the whole process by providing feedback control for each local layer (“always get ready”). Here we present a theoretical model to regularize voluntary motor control based on these two principles. The model features hierarchical organization and is composed of spiking building blocks based on the previous work in predictive coding and adaptive control theory. By simulating our manual interception paradigm, we show that the network could demonstrate motor preparation and execution, generate desired output trajectory following intention inputs, and exhibit comparable cortical and endpoint dynamics with the empirical data.

## 1. INTRODUCTION

Voluntary movement control is a complex process engaging many brain areas and multiple processing stages, typically including motor intention, preparation, and execution (see Kalaska 2009, Kalaska et al. 1997, Wise 1985, for review). How can actions be performed rapidly and efficiently through these stages? How exactly is motor intention transformed into action following these stages? Specifically, how this process is concretized within limited temporal and energetic resource? Recent advances in neuroscience have proposed many theories to account for these questions, e.g., representative view vs. dynamical system view in a broad sense of encoding and decoding (Georgopoulos et al. 1986, Shenoy et al. 2013), optimal subspace hypothesis for motor preparation (Churchland et al. 2010, Elsayed et al. 2016, Hennequin et al. 2014, Kaufman et al. 2014), theories of motor synergy for muscle generation (d’Avella et al. 2006, 2003), etc.

Yet it is still unclear how these theories can be related to give rise to the whole sensorimotor process from intention to execution. While the dynamical systems view, specifically the optimal subspace hypothesis (Churchland et al. 2010, Elsayed et al. 2016, Kaufman et al. 2014), provides insights into understanding the neural activity during motor preparation, its relation to the upstream motor intention, and downstream motor execution has not been fully addressed. Moreover, when and to what extent does a motor task need “motor preparation” is also intriguing. In the end, there is still a lack of a mechanistic account to bridge these phenomenological observations.

We argue that coherent mechanisms could underlie translation from motor intention to execution, to address this question by abstracting in twofolds. First, we see the process of motor intention → preparation → execution → final muscular distribution in the spinal cord, as an instance of gradual concretisation. In this regard, the existence of motor preparation lies in a linking stage to bridge the very broad “intention space” to the very specific “spinal execution space”, in a similar sense of the idea of “finding optimal subspace”. Backed by a multitude of training, the neural motor control system is like performing a narrow-down task in searching in the high-dimensional space by specialising the motor command through a hierarchical network, whereas supervised by the action feedback where necessary. Under certain pressure control, the stage of preparation can be either released promptly, or be constantly optimised by integrating incoming new information. In this high-dimensional space of reservoir of “search paths”, with more training, more alternative paths can be supplied for motor execution. Second, predictive coding has been shown to play an important role in extracting low-dimensional latent variables in a dynamical system (Recanatesi et al. 2021). As a general mechanism (Friston 2010), predictive coding can speed up this process by foreseeing the upstream information or by predicting next step based on the partial upstream information. As described in previous studies (Rao and Ballard 1999), the predictive processing mechanism achieves this by feedback control to each local layer to form internal models. In fact, recent work on monkey’s motor system also supports an “Always Prepare Hypothesis” (Ames et al. 2019), where predictive-flavoured neural activities are always engaged throughout the motor task, and run in parallel with motor execution. Altogether, by forming the “search paths” in the hierarchical high-dimensional motor space, and by ever predicting the outputs to the downstream hierarchy, these two principles can be summarised as “practice often” and “always get ready”.

Keeping these principles as guides to understand the motor control process, we need to specifically demonstrate that how can the neural system implement them. Neural motifs common in components but diverse in structure and functionality have been widely found in the neural system, such as functional columns in the visual cortices and other areas (Mountcastle 1997, Mountcastle et al. 1955, 1957). Although the structurally-similar columns have not been observed in the motor cortex and it is still under debate that whether such columnar structure has any functional significance (Horton and Adams 2005), the concept of motif can still play an important role in generating motor outputs, e.g., in terms of the theory of sub-movements (Crossman and Goodeve 1983, Elliott et al. 2001, 2017, Woodworth 1899). On the other hand, stable neural circuits prevail in the brain where balanced excitatory-inhibitory outputs are believed to be a key element to keep the brain without out of control (Denève and Machens 2016). It is therefore reasonable to postulate that the motor system is made up of stable building blocks as functionally-equivalent neural circuits. According to the above-described predictive coding doctrine, the internal dynamics of such building block should exhibit a predictive manner. Based on the previous work in predictive coding and adaptive control theory (Boerlin et al. 2013, Brendel et al. 2017, Denève et al. 2017), we constructed a spiking neural network that can both predict the next dynamics and behave as an autonomous system. This building block is built upon recurrent leaky-integrate-and-fire (LIF) spiking neurons and features tight excitation-inhibition (E-I) balance (through autoencoder architecture) commonly observed in the cortex. In addition, this building block can receive external feedback to update its dynamics, making it a trainable component in an error-based network. In this biologically-plausible manner, we come up with a homeostatic ensemble (termed homeostatic autoencoder unit, HAU) subserving as a basic functional unit to “always get ready”. See the methods section for the details on the HAU.

We then assembled a hierarchical model for motor control using the predictive building block HAU. It has been widely recognised that the neural network models generally utilise hierarchical structure and recurrent connections to mimic the biological system (LeCun et al. 2015, Sussillo and Abbott 2009). We follow this practice by constructing a two-layered hierarchical network. This network accepts motor intention and pressure control as inputs, and produce endpoint trajectory as output, with its layers explicitly demonstrating motor preparation/execution and spinal-muscle transformation. While the two layers represent motor-related cortices and spinal network respectively, they differ in anatomic functionality but share similar HAU structure. These two layers are uniquely defined by their location in the hierarchy and internal parameter settings. In this way, we explicitly construct this simple yet functional system using the building blocks, in an attempt to show that such a system is able to form a reservoir of “search paths” for motor control by sufficient training, i.e., by “practicing often”. See the methods section for the details on the network architecture and training methods.

In the end, we propose a theoretic framework to regularise voluntary motor control based on the principles of “practice often” and “always get ready”. Through simulating with our experimental paradigm of manual interception for monkey (Li et al. 2018), we show that the network can produce desired endpoint trajectory following intention inputs and can exhibit comparable cortical dynamics with the experimental data. We also show that the network underperforms without predictive power in the building block. These results suggest a coherent mechanism from motor intention to muscular execution, as well as provide a link to the related theories.

## 2. METHODS

### 2.1. Building block - HAU

#### 1. Structure

The HAU is a single-layered autoencoder driven by external inputs, error and predictive dynamics, based on the previous work of Boerlin et al. (2013), Brendel et al. (2017). The core of HAU, i.e., its internal layer, is composed of recurrently-connected LIF spiking neurons. Owing to the nature of LIF model, HAU is effectively a connected dynamical system, which in term leads to a general-purpose dynamics approximator. Figure 1 shows a scheme of an HAU, and its parameters are illustrated in Table 1.

**Table 1:** HAU parameter list

| Table 1: HAU parameter list |  |
| --- | --- |
| Parameter | Meaning |
| $\mathbf{c}, c_{0\sim 2}$ | input signals |
| $x$ | desired system state |
| $\dot{x}$ | derivative of desired system state |
| $\hat{x}, x_{0\sim 2}$ | estimated system states |
| $\dot{\hat{x}}$ | derivative of estimated system state |
| $\mathbf{F}$ | encoding connectivity |
| $\mathbf{D}$ | decoding connectivity |
| $r$ | LIF neuron's excitatory firing rates |
| $k$ | LIF neuron's excitatory spiking |
| $\Omega^e$ | excitatory recurrent connectivity |
| $\Omega^i$ | inhibitory recurrent connectivity |
| $\mathbf{K}$ | teaching positive error gain |
| $e$ | teaching error feedback |
| $f(x)$ | internal dynamics of the system state |
| $f(\hat{x})$ | internal dynamics of the estimated system state |
| $\lambda$ | leaky parameter |
| $\varphi()$ | nonlinear transformation |
| $V$ | LIF neuron's membrane potential |
| $\hat{V}$ | change of LIF neuron's membrane potential |
| $\mu, v$ | optimisation parameters |
| $\alpha$ | forward connectivity's learning parameter |
| $\beta$ | recurrent connectivity's learning parameter |

**FIG. 1:**
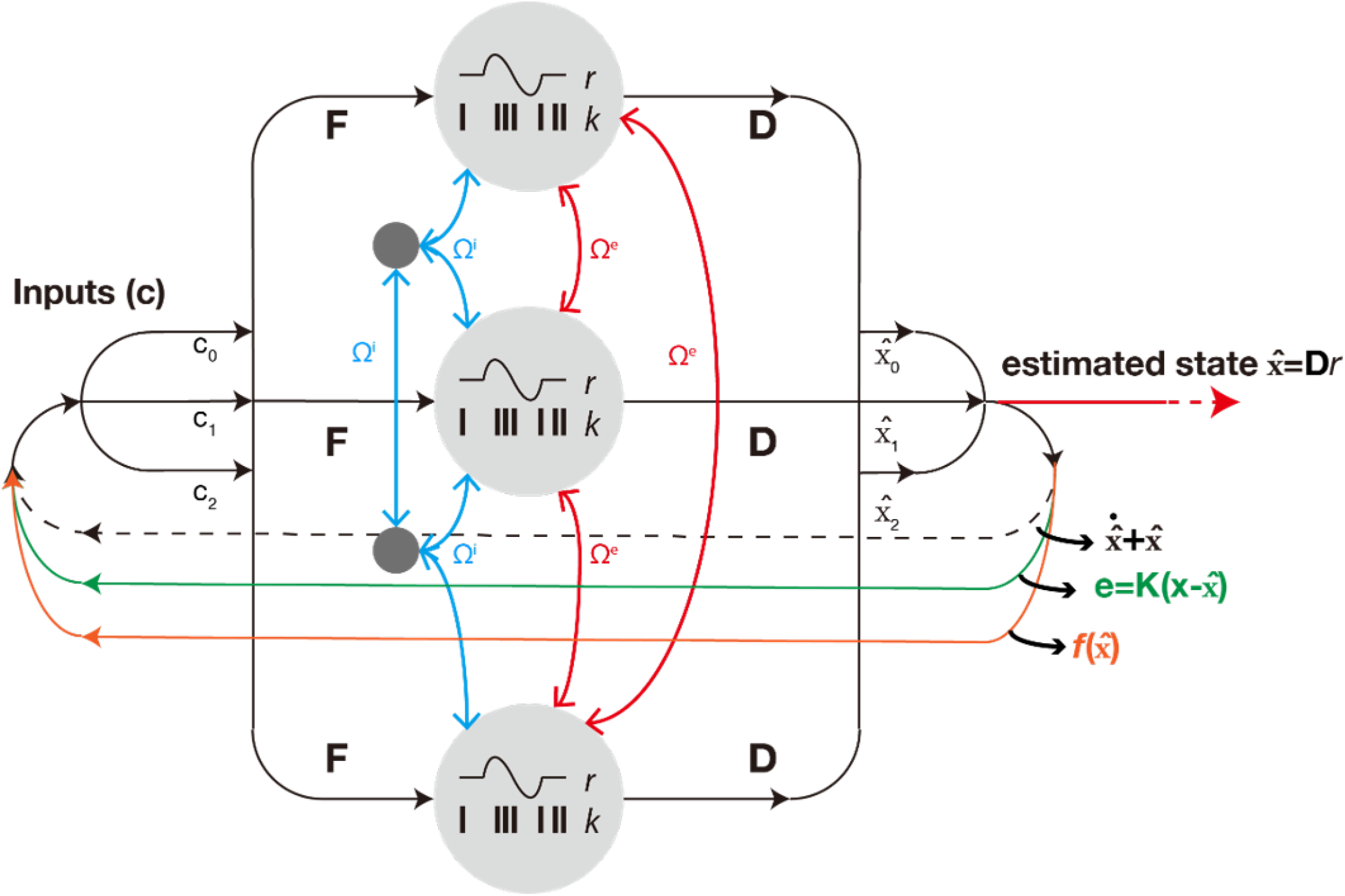
Scheme of HAU. The autoencoder approximates balance driven by both external and internal dynamics. Initially, HAU receives external inputs 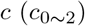 and treats the estimated internal state 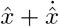 as a cancellation item to reach balance. Further, external teaching signal *e* and internal dynamics 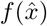 are added to the balance to force rich dynamics. The internal states are represented by the LIF model with **F** as forward connectivity and **Ω** as recurrent connectivity. The HAU is trained following the adaptive control theory (Boerlin et al. 2013, Brendel et al. 2017, Sanner and Slotine 1992, Slotine and Coetsee 1986). HAU can generate system state 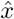 as outputs.

Suppose a dynamical system in the form of

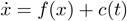

where HAU approximates the derivative as

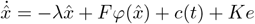

where *e*, as external error, is the difference between the desired and the estimated system states 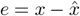

The dynamics of system state *x* then determines the change of the membrane potential *V* :

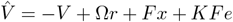

which follows the LIF model to generate spikes for any unit in the internal layer. As an autoencoder, HAU aims to optimise the loss function

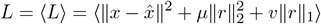

where *µ* and *v* are optimisation parameters.

One can see that an optimised E-I balanced membrane potential can reach the form of

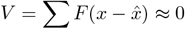

when, constrained by adaptive control theory (Boerlin et al. 2013, Brendel et al. 2017, Sanner and Slotine 1992, Slotine and Coetsee 1986), HAU converges as the effective inputs also approximate 0 and can reside in autonomous state.

According to Brendel et al. (2017), the updating rules of encoding connectivity is

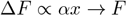

where the network learns to cover the input space; and the updating rule of recurrent connectivity is

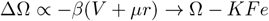

where network emulates a STDP-like mechanism.

One can also see that, due to the nature of autoencoder, HAU effectively forces the activities of the internal units to approximate zero, hence encouraging sparse representation. This makes HAU more biologically plausible.

#### 2. Adding predictive loop

We then make HAU even more powerful by adding predictive capabilities. As an autoencoder, its inputs boil down to subtracting the estimated output and adding the external error. So, we can rewrite the formula of effective input to the network as

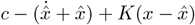

Meanwhile, *f* (·) is the core dynamics of HAU its internal layer. We can then add the prediction of its core dynamics 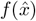 to the input to make it behave like an autoregressor

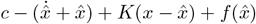

In this way, HAU now updates an internal estimate of state based on inputs. We can in turn see that, as a dynamic approximator, HAU can function as both an unsupervised autoencoder, driven primarily by inputs *c*, and as a supervised autoregressor, driven by inputs *c*, teaching signal 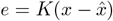 and internal dynamics prediction.

#### 3. Periodicity in the spinal cord

Previous studies have identified the mechanism of central pattern generator (CPG) of the spinal cord network (Hooper 2001). Research on spinal cord modelling develop the idea of CPG and point to a structure of rhythm generator (RG) + pattern formation (PF) + motorneurons (MNs) for the spinal cord (see Rybak et al. 2015, for review). In this work, we try to simplify these biomechanically-detailed approaches but adapt HAU to represent a functionally similar spinal cord network. Driven by motor command and a pre-defined periodicity generator, the HAU for spinal cord should form muscle patterns intrinsically and output velocity profile for a grouped muscle. We emphasise that this is obviously not a mechanistic modelling of the effectively much more sophisticated spinal cord network, but just a modulation and functionally similar nonlinear transformation of the motor commands.

### 2.2. Model architecture and training

In this work, we restrict the use of HAU to be layer-wise monolithic, i.e., in each layer of the hierarchical network there is only one HAU. This reduces the demand of computational power, but on the other hand, this also makes the simulation deterministic and less flexible.

It should be noted that, although HAU boils down to an autoencoder, the relation between its input and output is not like a “traditional” autoencoder. The HAU is designed to cancel “output” from input, but the “real” outputs are the estimated system state 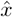

When the network goes beyond one layer, how to efficiently train it? For a deep neural network where its neurons are treated as single point emitting continuous values, powerful optimisation method like gradient-decent (GD) can be applied to train deep networks by error back-propagation. However, while the spiking neurons empower HAU to run in a biologically-plausible manner, it also casts problems to the efficient training. The discrete spiking output makes it impossible to use GD-like optimisation.

We adapt the difference target propagation (DTP) algorithm proposed in Lee et al. (2015) to train our HAU network. This algorithm attempts to emulate back-propagation via approximate inverse. Under HAU’s autoencoder setting, the inverse of the output can be straightforward. Then, using the proposed difference target propagation rule, we establish the feedback and forward mapping and send HAU’s internal states to the previous layers based on these mappings.

### 2.3. Data of simulation and neural recordings

Neural recordings data. We trained monkey B to perform manual interception (Li et al. 2018) task and used Blackrock Utah Arrays to record neural activities in M1 and PMd for each trial (Zhang et al. 2019). We analysed 508 correct trials of 42 neurons, and align them to the movement onset (for Figure 2B) and to the GO signal (for Figure 3). PCA was performed on these data averaging on the neurons. The first 3 dimensions of the reduced data were used as training and illustration purpose.

**FIG. 2:**
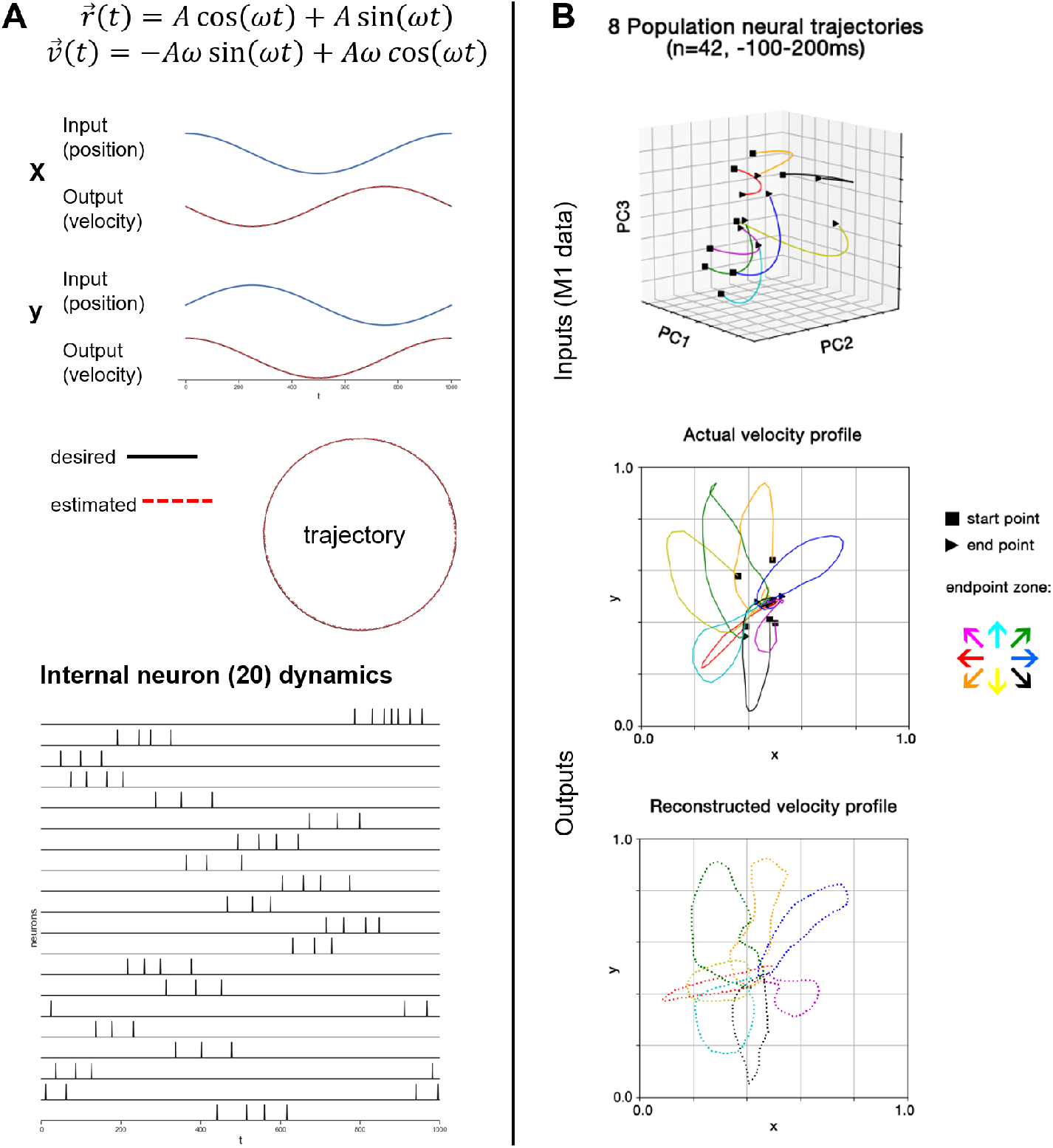
HAU simulating of a known dynamic of circular motion. **A**: HAU approximating known dynamics of circular motion. Inputs and outputs are both represented as 2-dimensional structures. With time progressing, the HAU receives positions as inputs and learns to output velocity. The desired trajectory follows the circular motion formula shown on the top. The spike trains of 20 internal LIF neurons are illustrated at the bottom. **B**: HAU approximating an unknown dynamics of spinal cord transformation (without periodicity modulation). For an interception paradigm, the 508 trials of 42 M1 neuron’s activities are averaged by neurons, and the first 3 dimensions of the PCA trajectory of the averaged data is shown on the top. For illustration purpose, only 8 trajectories are shown representing 8 endpoint zone directions. The HAU receives all 508 trajectories (motor commands) as inputs, and learn to output endpoint velocity profile. The actual and reconstructed velocity profiles are shown on the bottom.

**FIG. 3:**
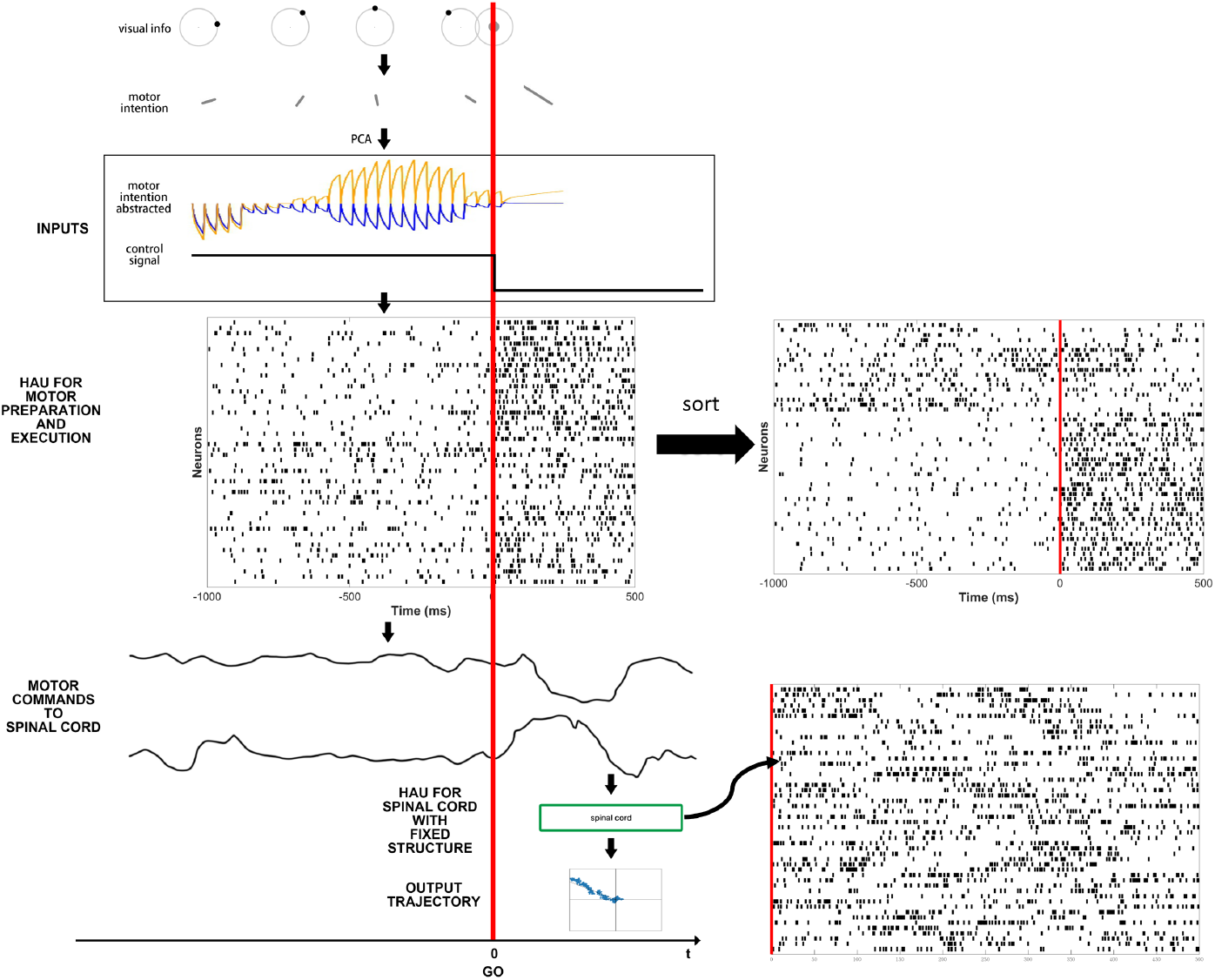
Two-layered HAU network simulating the motor process for manual interception task. **Top**: Inputs are generated from visual information and their correspondent motor intentions. See Methods section for the details. **Middle**: motor HAU receives abstracted motor intentions and control signal, and their internal dynamics show rich spike trains. A sorted version of this motor HAU on the right demonstrates a separation of neurons more oriented to motor execution and more oriented to motor preparation. **Bottom**: motor HAU outputs motor commands and spinal cord HAU with periodicity structure receives them to output a trajectory. Details of this spinal cord HAU on the right demonstrate traces of periodicity.

Simulation data. For the inputs in Figure 3, we simulated 1000ms of motor intentions. For a given interception endpoint zone, we generated the visual stimuli (screen) at each time bin (50ms). For each such stimuli, we generated a series of growing but brief movement endpoint trajectories to an advance angle, all represented as visual signal. Next, we used PCA to reduce these series of visual trajectories, and used first 2-dimensions as the inputs. This process is based on an ever-planning strategy for the interception task.

## 3. RESULTS

### 3.1. HAU simulation of both known and unknown dynamics

We first show the power of HAU in approximating dynamics. In a simple network, the performance of its composing element plays a dominant role in determining the network’s output. Therefore, before applying HAU in a more realistic motor control task, we need to demonstrate how HAU performs in simpler tasks.

Figure 2A shows the result of simulating a known circular motion. This one-layered network uses a HAU as its layer composed of 20 internal neurons. The network uses two-dimensional position as its inputs. During training, while trying to cancel the input position, the network is instructed to learn the velocity from known circular motion. Results in Figure 2A indicate that the network has been properly “taught” the dynamics and can output a circular trajectory, without being fed with the actual circular motion formula. The spikes of internal neurons reveal how does this network perform this task by its 20 neurons.

Figure 2B shows the result of simulating the unknown spinal transformation. This task is also performed on a one-layered network using a HAU as its layer, composed of 50 internal neurons. The network’s inputs are monkey’s M1 physiological neural data when performing manual interception task (see paradigm described in Li et al. 2018). We aim to output monkey’s manual velocity profile following these M1 commands. The activities of 42 M1 neuron during the 300ms around movement onset are reduced to 3-dimensional data using PCA, and in total 508 trials of such 3-dimensional data are used as motor command inputs. The explained variances plot indicates that this network using just 3-dimensional PCA input has been sufficient to represent the M1 dynamics. The result in Figure 2B shows that this one-layered HAU network has also learned how M1 commands can be translated through the sophisticated spinal cord network and can finally result into velocity. Sample of the reconstructed profile looks very similar with the actual one.

### 3.2. HAU network simulation of the entire motor control process of the manual interception paradigm

We then extend to a two-layered HAU network to simulate the neural process of motor intention → preparation/execution → spinal transformation. This process is based on our manual interception task for monkeys (Li et al. 2018). In this experiment, the animal is instructed to intercept a dot circularly moving around a circle. During each trial, the monkey’s hand is first fixed on the circle’s central point. Then a dot starts to rotate from a random position on the circle, at a certain speed and at a random direction (clockwise or counter-clockwise). After a GO signal, the monkey should immediately move its hand and try to intercept the still-moving dot. The monkey will get juice reward for successful interception. This paradigm is not complicated, but it has incorporated all the key elements in performing a motor task - prediction, intention, preparation, and execution. As shown in Figure 3, we set up a two-layered HAU network to simulate this process. As described in the Method, motor intention is represented as the PCA reduction of a series of visual moving paths. We use them as the inputs to the network. The first layer is composed of a HAU incorporating 50 neurons. We aim to show that this layer can translate the initial motor intention into motor commands fed to the spinal cord layer, so we term it as a general “motor HAU” without explicitly dividing into sub-areas of motor preparation, execution, plan selection, etc. The translated motor commands are then transmitted to the second layer with 50 internal neurons, where the pre-defined periodicity is applied. The output of this second layer is further reduced to two dimensions, corresponding to the integrated velocity profile, which is then finally plotted as endpoint trajectory.

We train this network using the difference target propagation algorithm described in the methods section. The network parameters in the model are carefully tuned to produce the trajectory in Figure 3, which looks very close to the initial input motor intentions.

Judging from the sorted version of the internal spike train of the motor HAU (middle right of Figure 3), one can clearly see the discrepancy of the neurons. Some neurons (bottom part) are much more dedicated to the execution section (after the GO signal), while some neurons (upper part) activate more balanced in both sections with a little busier before GO. One may argue that such discrepancy may arise primarily from the input patterns which employ an “ever-planning” strategy, but the heterogeneity of the motor HAU neurons is also comparable with the recent studies on the properties of motor-related cortical neurons, reminiscent of the well-observed M1 and PMd neural patterns studied in the previous literatures (Churchland and Shenoy 2007).

In addition, the internal spike train of the spinal cord HAU (bottom right of Figure 3) demonstrates periodicity, a kind of feature close to the EMG signals.

This result indicates that the HAU network can be applied in a complex hierarchical network. More importantly, it demonstrates that for a manual interception task, the HAU network can capture the core dynamics of motor-related areas and spinal cord network, and can simulate the process from motor intention to muscle output in its entirety.

### 3.3. Relation to the physiological data

While the two-layered HAU network can produce matched endpoint trajectory during the manual interception, it is unclear how its internal dynamics perform in relation to the biological neural system. One may argue that the initial input to the network, i.e., the reduced visual moving paths, is actually coined and hard to be measured in the biological system. Nonetheless, we can still check model’s relation to the monkey’s brain from its two HAU layers.

We compare both single neuron’s PSTH shapes and population trajectories between M1 and motor HAU, as shown in Figure 4. We choose two typical types of PSTH shapes from the 42 M1 neurons and then choose two closely matched from 50 motor HAU neurons. One can already see their similarity. To see a bigger picture of what is going on in general, we plot the PCA trajectories following the algorithm described in the Method. While the exact same structure cannot be observed in the motor HAU case, its trajectories clearly show the separation among 8 endpoint zone directions, and roughly traceable start to end dynamics, as comparable with the M1 case.

**FIG. 4:**
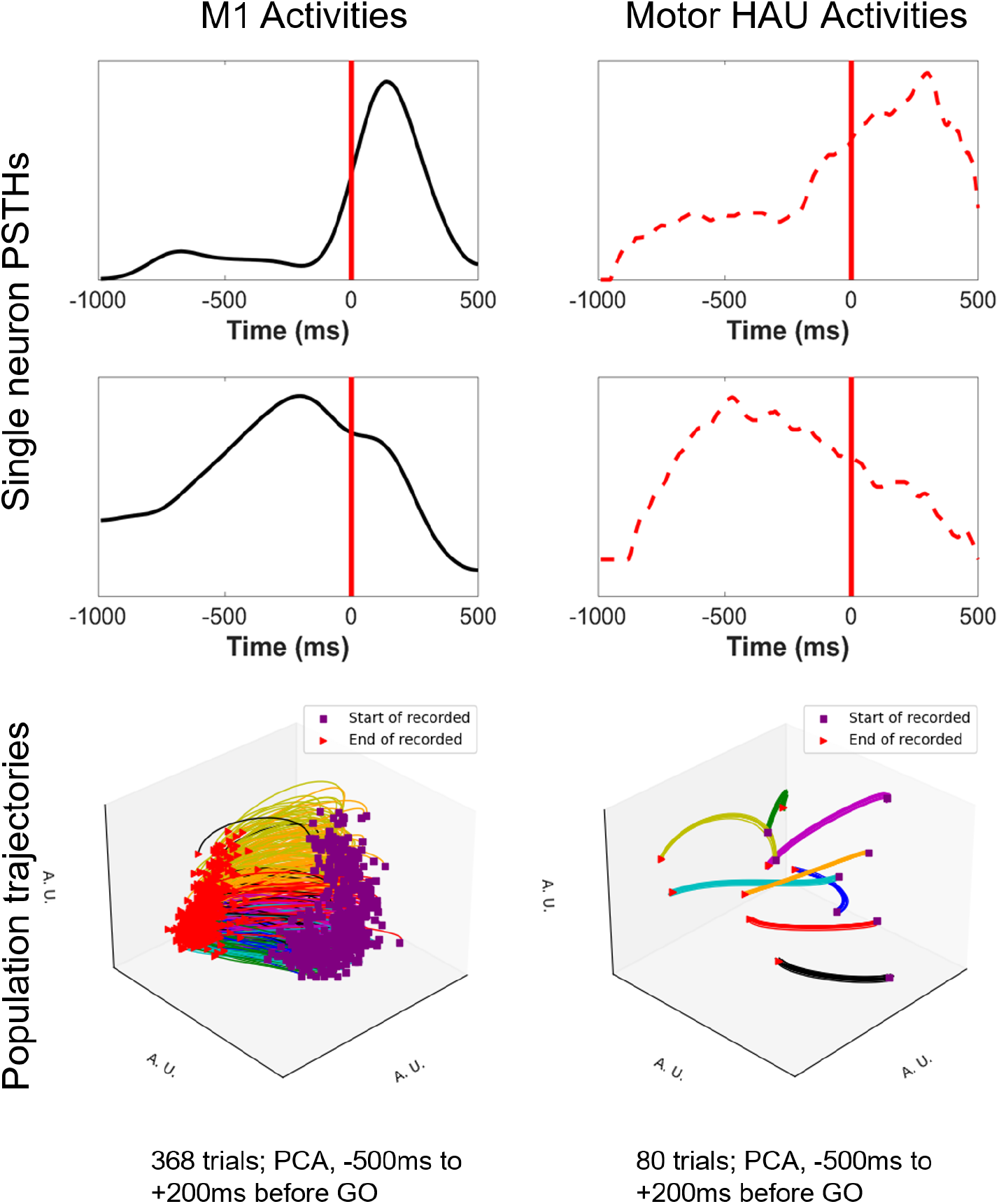
Comparison between M1 activities and motor HAU activities. **Left column**: Top: PSTHs of two sample neurons from the M1 neural recordings. Bottom: 504 trials average PCA of 42 M1 neurons. **Right column**: Top: PSTHs of two sample neurons from the motor HAU internal neurons, with similar PSTHs shapes to the correspondent M1 neurons. Bottom: 80 trials (10 for each direction) average PCA of 50 population internal neurons.

This correspondence between our model and our neural recording in M1 further illustrates that our model has captured the core dynamics of this motor task.

### 3.4. Results without predictive power in the building block

The above results show evidence to support the idea that by “practicing often”, the network can develop to a dynamical system underlying a motor task. Following an “ever-planning” strategy, the realistic features of the internal dynamics may shed light on the idea of “always getting ready”. But is the latter principle really necessary in this motor task?

We next consider how will the model perform without HAU’s predictive power. As described in the methods section, the “predictive loop” can be removed from the internal dynamics, which in turn transforms the HAU from an autoregressor into a supervised autoencoder. Using the same parameter settings and training epochs, we show in Figure 5 that, without the “predictive loop”, although the same network can still converge to the same results as the predictive case, it will struggle for a long period of training before finally reducing its error rate. The learning is clearly much slower without the predictive loop.

**FIG. 5:**
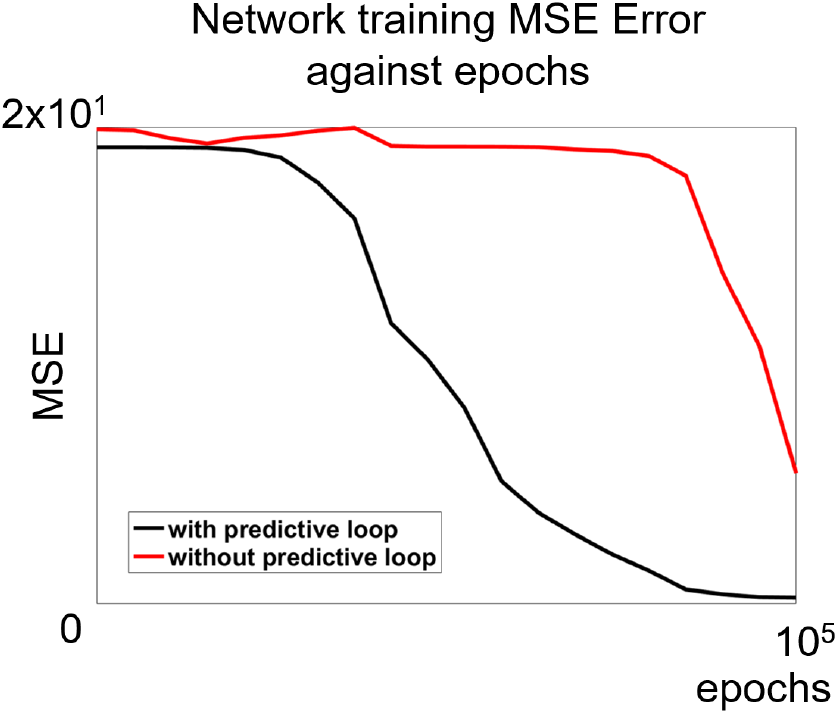
Difference with regard to predictive loop. The training MSE against epochs is shown for a typical training with and without predictive loop.

This result shows that the predictive nature of HAU network can speed-up the model’s convergence of generating a proper motor process. In the sense of a motor system, this may suggest that in order to perform a motor task efficiently one needs to “always get ready”.

## 4. DISCUSSION

In this paper, we show a mechanistic model for voluntary motor control that can produce desired trajectory according to the input motor intention. This model is made upon predictive building blocks, HAUs, that assemble a hierarchical structure to allow a discrete gradient of transformations.

When performing a motor task, this hierarchical predictive model transforms the initial intention into actions feedforwardly. Owing to the delayed control signal, discrepancy of neurons for motor preparation and execution emerges during the course. We emphasize two essential elements underlying the modelling work: “practice often” and “always get ready”. The former refers to the network training with the hierarchical structure, while the latter is realised by predictive HAUs in each network layer. We show that our modelling results generate comparable neural population statistics with the physiological data, and therefore we conclude that our model featuring these two characteristics can provide insights into understanding voluntary motor control for certain experimental tasks.

### 4.1. Significance of the work

We see the use of predictive building blocks of hierarchical spiking neurons to simulate the whole motor control process as the essential contributions of our work, and we believe it is significant to understand the motor system with simple rules of predictive coding and hierarchical training.

Predictive coding has become one of the important hypotheses of the underlying neural mechanisms. In the recent years, it has gained many experimental supports (Millidge et al. 2021), especially in the sensory systems. We consider that the predictive capabilities serve as an essential and crucial enterprise of performing any motor tasks, and with this regard the neural substrates governing motor system should also exhibit predictive traits. The HAU with built-in predictive power thus resides at the centre of the model as “predictive populations”. Essentially, we found in Figure 5 that the system may work but in a much slower manner without predictive power, supportive to the idea that predictive capabilities are crucial for the motor system. Our “always get ready” principle can potentially account for the highly-innervated neural substrates in the motor-related cortices for motor preparation and execution, and such accountability is consistent with the recent founding of simultaneous motor preparation and execution (Ames et al. 2019), where an “always-on” predictive mechanism is vert plausible.

We then show that the network can complete a simple motor task by cascading the HAUs hierarchically. In the meantime, it is worth noted that although the HAUs communicate with each other using continuous signals, their internal computations are effectively performed by spiking neurons. These as a whole make the entire model a hierarchical spiking neural network at its core, where predictive populations transform signals at each layer. We consider our model biologically-plausible at the population level.

### 4.2. Are we claiming an anatomically equivalent implementation?

While the cortical pathway of motor system has been illustrated in detail in the previous literatures (see Strick et al. 2021, for a recent review), the specific contributions of the cortical and subcortical areas are still largely unclear. Typically, in a delayed task, the arising, development and causality of motor intention, preparation and execution essentially involve distributed processing and multi-region coordination. Yet, how these interactions are realised at the level of neural circuits or biophysics are highly debated. This work attempts to provide an insight into understanding this process in a theoretical model, and aims only to propose possible mechanistic guidance that can lead to the transformation from motor intention into execution. Indeed, what we propose in this work is how this transformation can occur functionally and conceptually, but not claiming an anatomically equivalent neural implementation. Based on the interception paradigm, we show that heterogeneous neurons dedicated for motor preparation and execution can emerge in a single HAU through training, but the actual functioning body in the neural system may be distributed in multiple regions. On the implications of our model, we are proposing a modular architecture through HAUs at the population level, but not necessarily a modular or columnar mapping in the anatomical structure. Likewise, we are not proposing that the neural system is implementing autoencoder or alike structure in any way. Instead, we advocate that the voluntary motor control in general can be better framed and abstracted beyond the variable single neurons at the population level, where gradual specialisation and predictive processing can be seen as guiding principles to bridge the “microscopic” neuronal activities and “macroscopical” behaviours.

### 4.3. On dynamical system and optimal feedback control

The view of dynamical system has played an important role in understanding motor control in the recent years. The related work typically uses dimension reduction methods to transform the population neural activities into low-dimensional state space, where the development and interaction of neural activities can be traced and analysed (Churchland et al. 2010, Kaufman et al. 2014, Shenoy et al. 2013). In line with such approaches, our work, as shown in Figure 4, also demonstrates such state space traces for each trial, comparable with the experimental results. Moreover, recent work also showed a general picture of how predictive coding mechanisms could give rise to low-dimensional representations. In this sense, our work present consistent results for the motor system, though further investigations may be worth on studying the underlying connections. In addition, more recently, there has been a focus on how feedback control, especially optimal feedback control (OFC) theory (Todorov 2000, Todorov and I. 2002), can serve as an intrinsic mechanism of the dynamical system to further promote the control process (Kao et al. 2021). Our model does not explicitly implement OFC as an internal mechanism, but do includes feedback loops to regularise the model owing to the nature of autoencoder structure. Although feedback is already a built-in mechanism for the predictive HAU, however, in the network level, we explicitly omit the feedback loops that can possibly integrate external information (sensory or proprioceptive) and back-propagate to the motor system. This omission is largely due to the feedforward nature of the interception paradigm, where motor plan should be enough to lead to successful task without further adjustment during the rapid course of motor execution (Li et al. 2018).

### 4.4. Future directions

Our work develops a parsimonious approach to account for the transformation of motor intention into motor action, based on relatively simple rules of predictive coding and hierarchical training. Our results show that, given a simple interception task, these rules suffice at guiding the motor system to fulfil the whole process in a feedforward-only manner. Beyond this very-simplified scenario, we see further investigations may be worth in the following directions based on these two rules. First, to make the model more realistic and applicable to the more complex tasks, explicit feedback loop in the network level should be added to form a more powerful motor control framework. Besides, the optimal feedback control mechanism may be also necessary in the building block level. Having an obviously more complex network, what follows is a better training method for the multi-layered SNNs. Recent advances have been focused on new approaches to optimise the training and tuning of big SNNs (see Taherkhani et al. 2020, for a recent review), with which our model, equipped with feedback structures, may generate much more flexible control strategy and suit multi-modal inputs. Having a better training method at hand, one may also try to introduce paralleled building blocks to extend the network laterally, which could potentially improve the overall robustness of the model. In the end, biological-inspired methods such as explicit internal models at the building-block level, reservoir of submovements, etc. could be considered to the model towards a more flexible and robust motor control system.

